# Complete mitochondrial genome of *Muricea crassa* and *Muricea purpurea* (Anthozoa: Octocorallia) from the eastern tropical Pacific

**DOI:** 10.1101/042945

**Authors:** Angelo Poliseno, Odalisca Breedy, Michael Eitel, Gert Wöerheide, Hector M. Guzman, Stefan Krebs, Helmuth Blum, Sergio Vargas

**Affiliations:** Department of Earth and Environmental Sciences, Palaeontology & Geobiology, Ludwig-Maximilians-Universität München, Richard-Wagner-Straße 10, 80333 München, Germany; Centro de Investigación en Estructuras Microscópicas, Universidad de Costa Rica. P.O. Box 11501-2060, Universidad de Costa Rica, San José, Costa Rica; Centro de Investigacin en Ciencias del Mar y Limnología, Escuela de Biología, Universidad de Costa Rica. P.O. Box 11501-2060, Universidad de Costa Rica, San José, Costa Rica; Smithsonian Tropical Research Institute, P.O. Box 0843-03092 Panama, Panama; GeoBio-Center, Ludwig-Maximilians-Universität München, Richard-Wagner-Straße 10, 80333 München, Germany; Bayerische Staatssammlung für Paläontologie und Geologie, Richard-Wagner-Straße 10, 80333 München, Germany; Laboratory for Functional Genome Analysis (LAFUGA), Gene Center, Ludwig-Maximilians-Universität München, Feodor-Lynen-Strasse 25, 81377 München, Germany

**Keywords:** mitogenome, *Muricea crassa*, *Muricea purpurea*, NGS

## Abstract

We sequenced the complete mitogenomes of two eastern tropical Pacific gorgonians, *Muricea crassa* and *Muricea purpurea,* using NGS technologies. The assembled mitogenomes of *M. crassa* and *M. purpurea* were 19,586 bp and 19,358 bp in length, with a GC-content ranging from 36.0% to 36.1%, respectively. The two mitogenomes had the same gene arrangement consisting of 14 protein-coding genes, two rRNAs and one tRNA. Mitogenome identity was 98.5%. The intergenic regions between *COB* and *NAD6* and between *NAD5* and *NAD4* were polymorphic in length with a high level of nucleotide diversity. Based on a concatenated dataset of 14 mitochondrial protein-coding genes we inferred the phylogeny of 26 octocoral species.

---

*Muricea crassa* (Verrill, 1869) and *Muricea purpurea* (Verrill, 1864) are two shallow water gorgonians of the family Plexauridae. Their distribution is limited to the eastern tropical Pacific where they are abundant members of coral communities and littoral zones (Guzman et al., 2004). Samples were collected as part of an ecological and biodiversity survey undertaken in the Coiba National Park (Panama). Genomic DNA was extracted from ethanol-preserved samples and was used to construct genomic libraries using the Accel-NGS 1S DNA Library kit (Swift Biosciences, Ann Arbor, MI, USA) following the manufacturers instructions. These libraries were sequenced (100bp PE) on an Illumina HiSeq (Illumina Inc., San Diego, CA). The quality of the reads obtained was assessed with FastQC (Andrews, 2010), low quality reads and Illumina adaptors were trimmed using Trimmomatic 0.3.2 called from Trinity RNA-Seq 2.0.6 (Grabherr et al., 2011). Despite its original purpose, the Trinity RNA-Seq assembler was used after normalization to 50X coverage for *de-novo* mitogenome assembly. The assembly resulted in a single mitochondrial contig in both species. Initial annotation was performed with the ORF finder function implemented in Geneious 8.1.7 (Kearse et al., 2012) and was corroborated by comparison with published octocoral mitogenomes. The presence of DNA repeats was assessed with the tandem repeats finder server 4.08 available at https://tandem.bu.edu/trf/trf.html (Benson, 1999). The complete mitogenomes of *M. crassa* (LT174652) and *M. purpurea* (LT174653) were 19,586 bp and 19,358 bp long, with a GCcontent of 36.0% (*M. purpurea)* and 36.1% (*M. crassa*), respectively. Both mitogenomes had gene arrangement of type “A” (see Brockman and McFadden, 2012). In total, the Coding DNA Sequences (CDSs) spanned about 76% of the mitogenome in both species. Among protein-coding genes, the highest level of nucleotide diversity (0.4%) was found in *NAD1*, *NAD6* and *COX2*, whereas no nucleotide substitutions were found in *NAD3, ATP6* and *ATP8*. Except for *NAD2* and *NAD5* (13bp overlap), the other protein-coding genes were separated by intergenic regions (IGRs) of different lengths. In both species, the shortest IGRs were those located between 12S rRNA and *NAD1* and between 16S rRNA and *NAD2*, while the longest was found between *NAD5* and *NAD4*. The latter IGR was also the most diverse region with a nucleotide diversity of 6.2%. Length polymorphism was found in the *COB-NAD6* IGR, which was 184bp shorter in *M. purpurea* than in *M. crassa*. Sequencing of indel-rich IGRs such as that between *COB* and *NAD6* may result useful for molecular species-identification in the genus *Muricea*. Finally, we found a 37 bp tandem repeat in the IGR between *NAD4* and tRNA of *M. crassa*.

The two newly sequenced complete mitogenomes were used to assess the phylogenetic relationships among 26 different octocoral species. A concatenated nucleotide alignment of 14 protein-coding genes (15,249 bp in total) for 41 taxa was generated with MUSCLE (Edgar, 2004) using the default options provided in Seaview (Gouy et al., 2010). The maximum likelihood tree was inferred in RAxML 7.2.8 (Stamatakis, 2006) under the GTR+GAMMA substitution model. Node support was estimated using 1000 bootstrap pseudoreplicates. The phylogenetic tree (Figure 1) was re-rooted using three calcaxonians and two pennatulaceans as outgroup (not shown in Figure 1). Tree topology is consistent with recently published studies (Figueroa and Baco, 2015). The phylogenetic placement of *Muricea* spp. sister to Nephtheidae (*Dendronephthya* spp. and *Scleronephthya* spp.) is likely an artifact caused by poor taxon sampling in the family Plexauridae.

**Figure 1.**
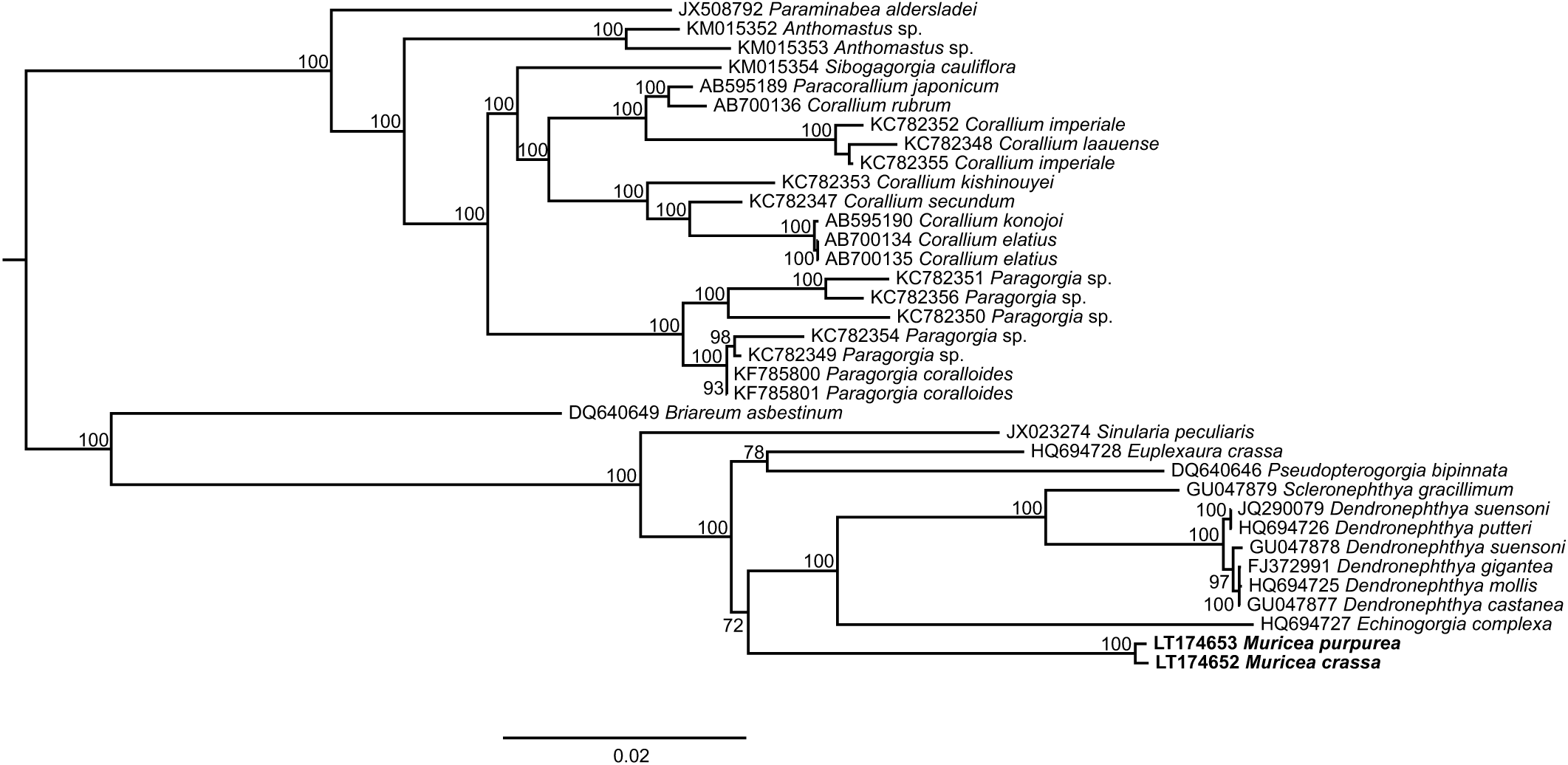
Phylogenetic tree of 41 octocorals based on a concatenated alignment of 14 mitochondrial protein-coding genes. The calcaxonians (Keratoisidinae sp*.,Acanella ebumea, Narella hawaiinensis* and *Junceella fragilis)* and pennatulaceans (*Renilla inuelleri* and *Stylatula elongata*) were used to re-root the tree but are not shown here. Numbers at the nodes indicate bootstrap values.

## Acknowledgements

We would like to thank the Gene Center (Ludwig-Maximilians-Universität, München) for technical support and sequencing. SV is indebted to N. Villalobos, M. Vargas and S. Vargas for their constant support.

